# Analysis of intracellular transport dynamics using quantitative phase imaging and FRET-based calcium sensors

**DOI:** 10.1101/2023.05.05.539565

**Authors:** Robert E Highland, Albert Rancu, Hillel Price, Steven M Parker, Meghan Reynolds, Brenton D. Hoffman, Adam Wax

## Abstract

Understanding cellular responses to mechanical environmental stimuli is an essential goal of cellular mechanotransduction studies and remains a target for microscopy technique development. While fluorescence microscopy has been advanced for this purpose - its molecular sensitivity, the ability of quantitative phase imaging to visualize subcellular structure has yet to be widely applied, perhaps due to its limited specificity. Here we seek to combine Quantitative Phase Imaging (QPI) with a molecularly sensitive Förster resonance energy transfer (FRET) construct for cell mechanotransduction studies. The multimodal imaging instrument is applied to examine cellular response to hypo-osmotic stimulus by observing the influx of calcium ions using a FRET based sensor coupled with mapping of the redistribution of intracellular mass using QPI. The combined imaging modality enables discrimination of cell response by localized region and reveals distinct behavior for each. The analysis shows cell flattening and oscillatory mass transport in response to the stimulus. With the broad array of FRET sensors under development, the combination with QPI offers new avenues for studying cell response to environmental stimuli.

## Introduction

Interplay between cells and the dynamic microenvironment occurs through mechanical, electrical, and chemical mechanisms [1]. Environmental cues can induce a range of changes in cells’ internal state. For example, cells can respond to changes in their environment by altering the transmission state of an ion channel, allowing for active ion exchange with the microenvironment [2–4]. Fluorescence imaging has been the premier imaging tool in cellular biology to observe cell responses to microenvironment perturbations by visualizing both structural and electrochemical dynamics at high resolutions [5–8]. Förster resonance energy transfer (FRET) is a nanoscale sensing variant of fluorescent imaging where an excited donor fluorophore non-radiatively transfers energy to an acceptor fluorophore to measure their proximity [9]. FRET can be implemented with a wide variety of fluorophores coupled to molecules of interest, creating intracellular sensors, to uniquely report on cellular changes [10–12]. FRET sensor engineering allows for the modality to capture molecular events, though the fluorescence-based approach is limited in terms of observing cellular morphological changes without additional labeling. Additionally, the technique can be restrictive due to the need for long exposure times which must also be balanced against photobleaching of the sensors and subsequent signal loss. This results in poor temporal resolution and limited capabilities for continually observing dynamic processes [13].

Quantitative Phase Imaging (QPI) presents an alternative which can avoid some of these shortcomings. QPI measures the phase delay of light retarded by a thin, weakly scattering sample and uses this as a label-free contrast mechanism. Consequently, this imaging modality does not rely on exogenous fluorophores within the cell and therefore avoids the negative effects of photobleaching in FRET, allowing for a higher temporal resolution and long, continual exposures. Moreover, the quantitative nature of the modality does not depend on the intensity of light collected and facilitates the use of an array of analysis methodologies for examining biological cell processes. QPI yields information on the mass distribution within the cell using optical volume (OV), a measure related to the dry mass of the cell by a scaling constant [14]. OV has been shown to be a useful marker of morphological changes and as a diagnostic indicator for blood cells [14–19]. Advantages of QPI such as increased temporal sampling, longer duration studies and whole cell mass distribution imaging suggests its utility in supplementing the FRET modality[19–21].

Here we present a novel QPI-guided FRET imaging technique and its application in measuring cell physiological reaction following a hypo-osmotic stimulus. This technique builds on previous work showing the ability to combine the two modalities into a single optical system [22]. However, this report illustrates how information obtained simultaneously by both systems can be used together to further understand cellular responses to stress. Osmotic stimulus is applied to A431 skin cancer cells transfected to express the Twitch-3 calcium ion-binding FRET sensor [12], and continuous cellular response is recorded simultaneously for both modalities. QPI information is used to delineate approximate cell morphology and analyzed to create an automated subcellular segmentation algorithm, providing spatial discrimination for analyzing region-specific activity as measured by OV. The QPI information is registered with FRET images, enabling FRET information to be analyzed by cellular region. Thus, the combination of modalities enables long time analysis of calcium ion dynamics and the influence on intracellular mass dynamics, while circumventing pitfalls associated with intensity-based FRET images.

## Materials and Methods

### Combined QPI-FRET System

The QPI-FRET imaging system used throughout experimentation was designed to capture QPM via digital holography and FRET readings, simultaneously (Fig 1). A former iteration of the QPI-FRET system has been described and used to visualize intracellular morphological changes and apoptotic activity [22]. The QPI light source is a 660 nm diode laser deployed in a previously described Mach-Zehnder off-axis interferometry scheme. Light from the QPI source is divided into sample and reference arms using a 90/10 fiber optic splitter. Light from the sample arm is passed through a sample on a heated stage and collected using a Nikon 100×, 0.9 NA objective. Sample arm light is then recombined with the reference arm light using a 50/50 beamsplitter (BS) and projected to a CCD + CMOS camera (Point Grey: Grasshopper3, Richmond, BC, Canada).

**Fig 1.**
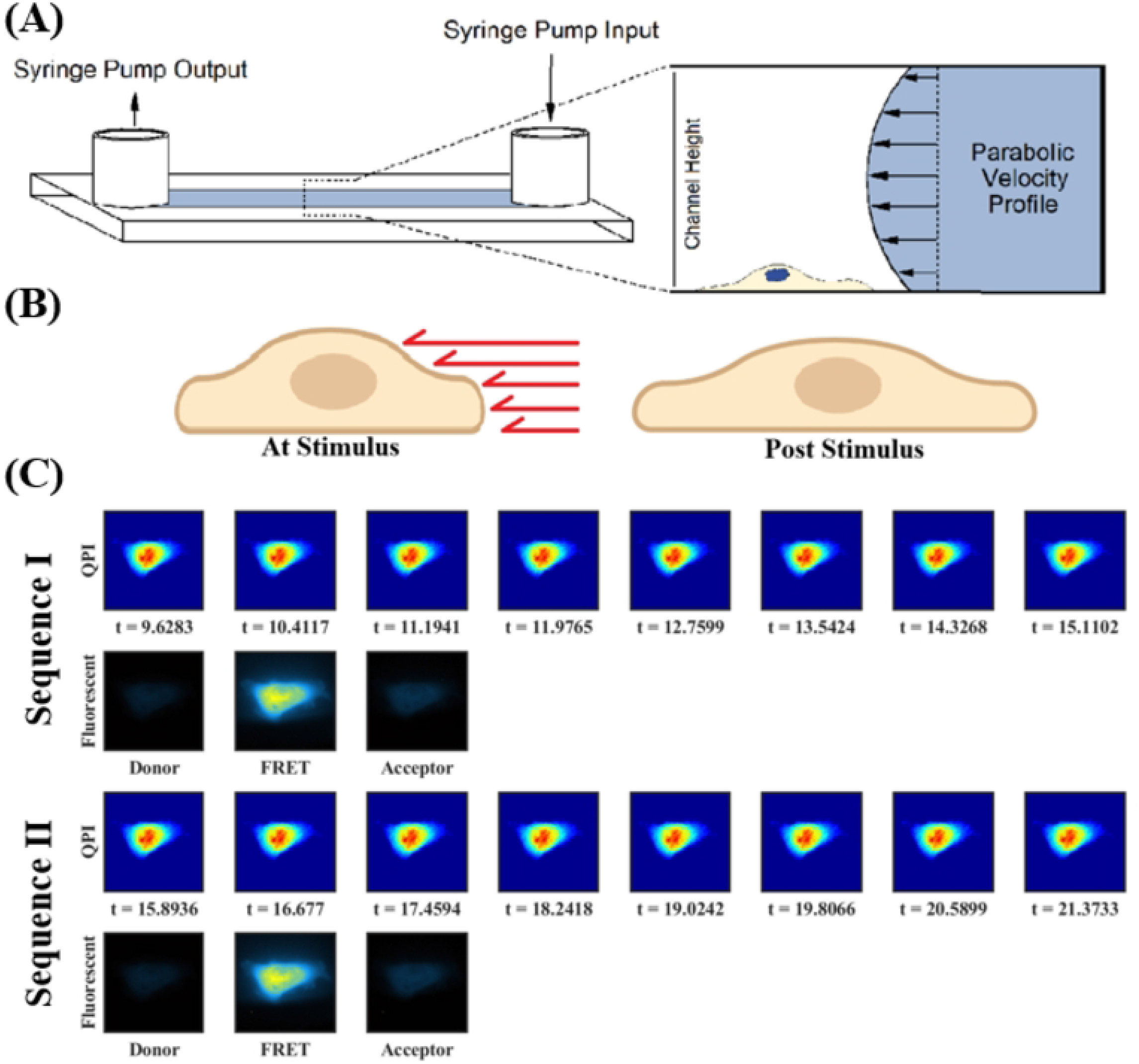
Combined QPI-FRET System. (A) The optical system diagram for simultaneous quantitative phase microscopy and fluorescent imaging. The FRET excitation, FRET emission, and QPI imaging wavelengths are spectrally separated using sets of dichroic mirrors (DCM) and fluorescent filter wheels. (B) Spectra for excitation filters (EX), emission filter (EM) and dichroic mirror (DCM).

The filter set used to detect ECFP-cpCitrine174 FRET sensors included, ECFP excitation (ET440/40×) [EX1], ECFP emission (ET473/24m) [EM1], cCitrine174 excitation (ZET514/10×) [EX2], cpCirine174 emission (ZET560/80m) [EM2], and a dichroic mirror (ZT442/514rpc) [DCM]. (Fig 2) FRET detection was accomplished by using three fluorescence exposures, acquiring images in the donor channel (ECFP excitation, ECFP emission), the FRET channel (ECFP excitation, cpCitrine174 emission), and the acceptor channel (cpCitrine174 excitation, cpCitrine174 emission) each with the same exposure time (∼800 ms). In addition to utilizing this relatively short exposure time, further fluorescence photobleaching was limited by blocking subsequent excitation light via an electronically controlled shutter in successive sequences. The FRET ratio was calculated as the FRET emission divided by the donor emission.

**Fig 2.**
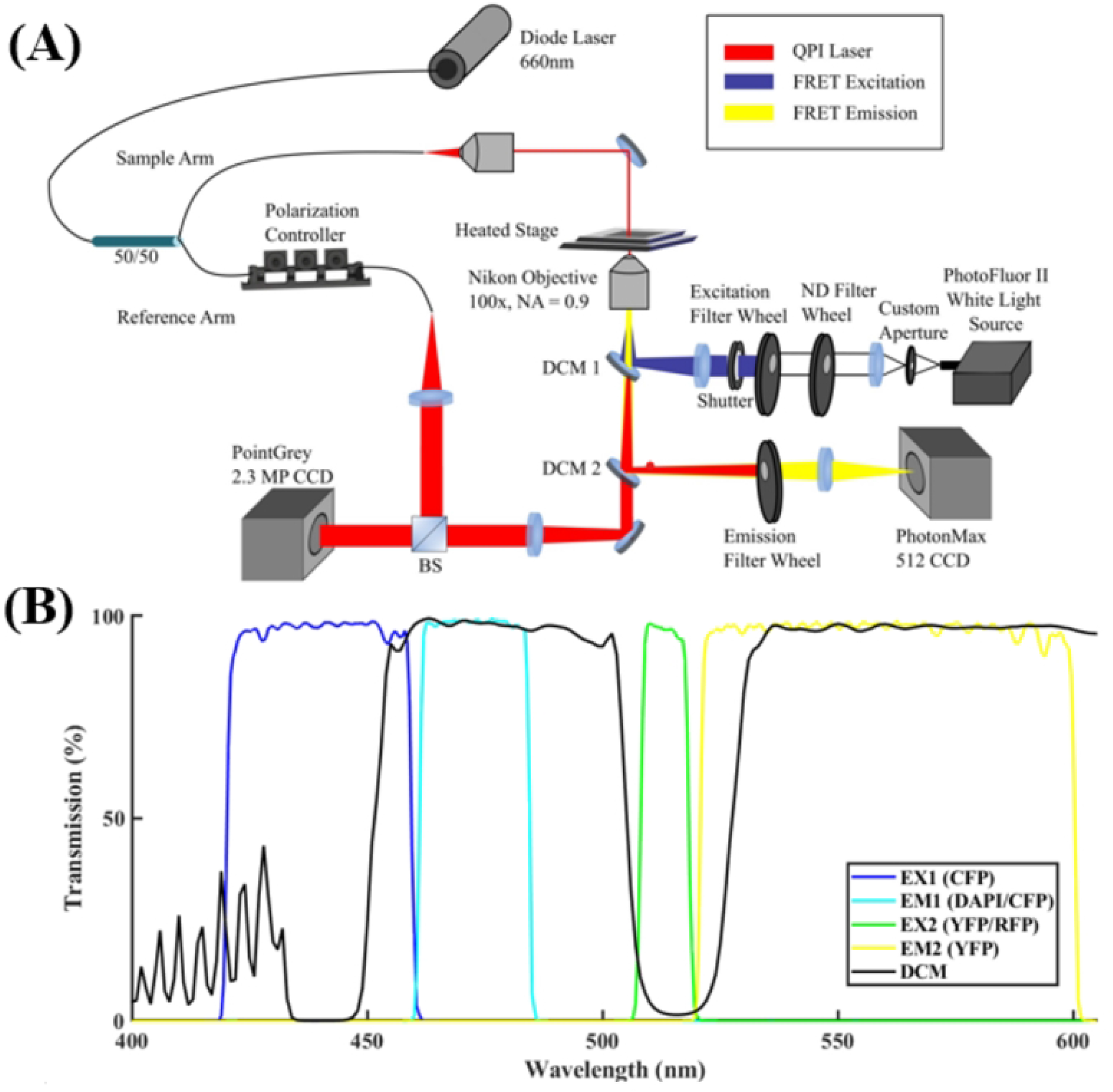
Experimental Setup and Sample Imaging Acquisition. (A) Depiction of flow channel with parabolic flow. Channel geometry and fluid viscosity define cellular shear stress. (B) Expected cell behavior upon stimulus, red arrows indicate relative shear stress at varying cell height. (C) Simultaneous QPI-fluorescent imaging workflow over two imaging sequences. QPI images are acquired every 0.815 s with FRET sequences interleaved simultaneously during the first three QPI frames.

### Cell Culture and Transfection

A431 epithelial carcinoma cells were cultured at 37°C in Dulbecco’s Modified Eagle’s Medium supplemented with 10% fetal bovine serum and 1% Pen-Strep antibiotic in a 5% CO_2_ environment. The cell line was passaged every 2 days using 0.25% trypsin at approximately 75% confluency. In preparation for transient transfection, A431 cells were plated in a single well of a six-well plate at 60% confluency for approximately 30 hours. Cells were washed with PBS and existing media was replaced with antibiotic-free media. Transient transfection was then performed using Lipofectamine 2000, following manufacturer protocol. A431 cells were transfected with pcDNA3 Twitch-3, a fluorescent reporter for calcium signaling (2 ng of plasmid DNA). The Genetically Encoded Calcium Indicator (GECI), Twitch-3, functions via a modified Troponin C (TNC) domain which binds, limitedly, to Ca^2+^ to enable ratiometric FRET sensing[12]. Two days following transfection, cells were seeded in fibronectin-coated glass chambered flow slides (Ibidi 0.4 μ-Slides) in preparation for imaging (Fig 2A).

### Cell imaging protocol

Because the growth media produced auto fluorescence, it was replaced immediately prior to the imaging session with Live Cell Imaging Solution + 5% FBS and allowed to incubate for 10 minutes. Following the incubation period, the chambered flow slide was placed onto an Ibidi stage heater (37°C) to maintain thermal equilibrium. Hypo-osmotic fluid stimulus (1:1 Live Cell Imaging Solution and Deionized Water) was applied to A431 cells intermittently during flow using a NE-4000 Two Channel Programmable Syringe Pump. Initially, cells were exposed to imaging media for 28 seconds, immediately followed by a hypo-osmotic shock for 5 seconds at a rate of 6 mL/min, followed by further exposure to imaging media (Fig 2B).

The imaging sequence used throughout experimentation is shown in Fig 2C. Each imaging sequence comprises eight QPI frames, acquired every 0.8 s and three fluorescent frames, which coincide with the first three QPI frames. QPI acquisitions can occur simultaneously with FRET since the wavelength used is spectrally distinct from the fluorescence emissions used for FRET imaging. The final five frames of the sequence includes only QPI imaging.

### QPI Image Segmentation

Automated QPI Segmentation is introduced here as a novel method for visualizing changes in mass distribution throughout a cell using QPI. The segmentation algorithm defines three physiologically relevant “layers” of the cell based on the Optical Volume (OV). Based on typical subcellular morphology of the cell lines, the inner layer corresponds to the most optically thick part of the cell, and likely containing the nucleus [22], a middle layer that is expected to contain the endomembrane system, specifically the endoplasmic reticulum, where calcium regulation occurs, and lastly an outer layer, where cellular actin reorganization is hypothesized to occur after stimulus. Fig 3 demonstrates the application of this segmentation to a representative cell in a single frame of QPI data acquired before application of osmotic stimulus. Segmentation boundary conditions were calculated dynamically on a per-frame basis to eliminate bias that may occur when the average phase of the cell increases or decreases during the experiment. QPI images of cells were masked above a threshold of 0.25 radians, representing the boundary between the outer layer and the extracellular environment. The mean phase of the cell body was measured and used as the second threshold for the boundary between middle and outer layer. The inner layer was chosen as all phase values higher than 1.5 times the mean phase of the cell. The OV of each layer was determined and monitored across the time course of each experiment. Cell layers were nominally elliptical and any holes in each layer where individual pixel values fell below the respective QPI threshold were accounted for and included in the proper layer volumetric measurement to ensure consistent integrity of each region across the duration of the experiment.

**Fig 3.**
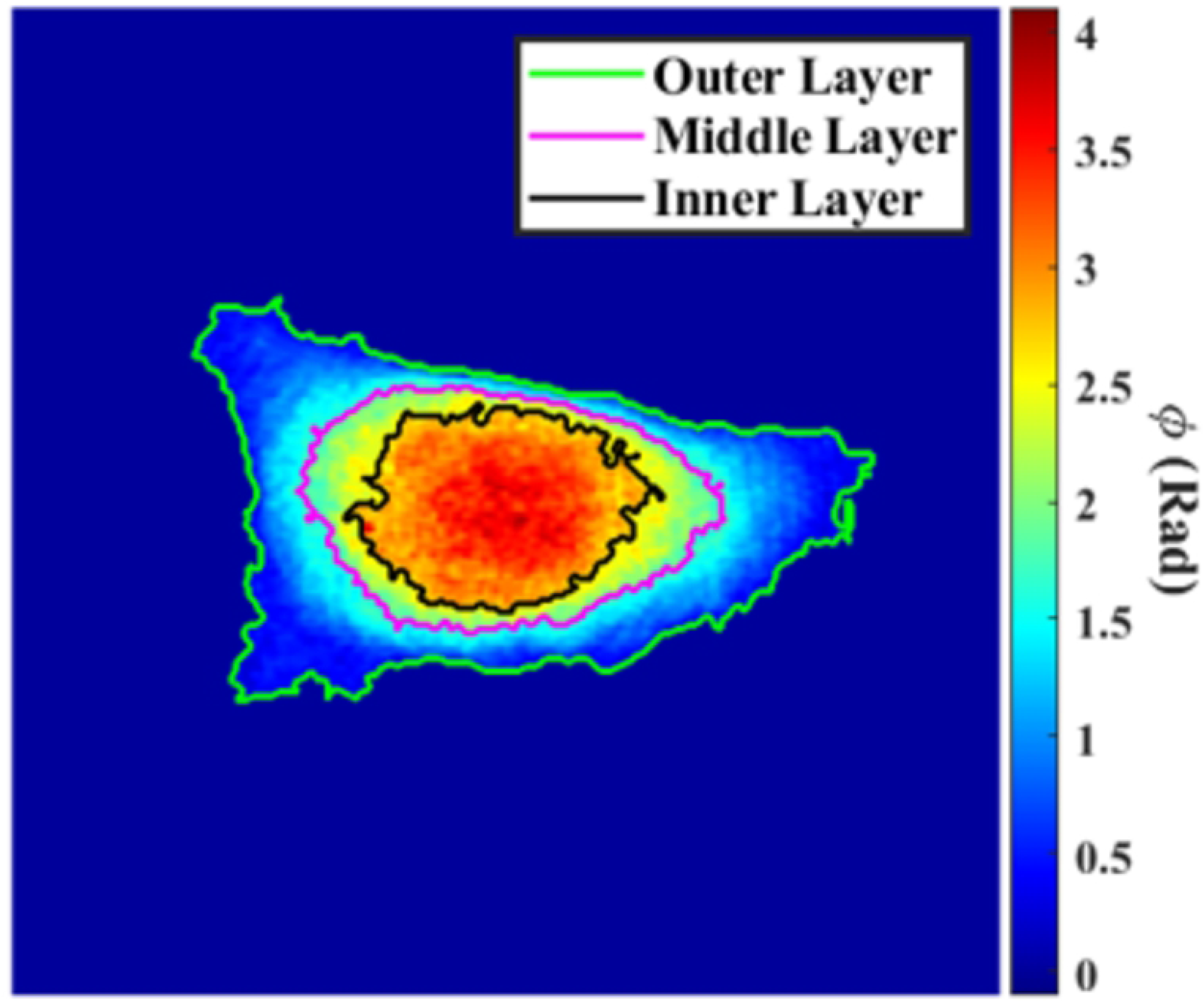
Example QPI image with automated segmentation layers applied.

### Optical Volume

Optical Volume was measured from phase images using previously described methods [14–18]. Optical path length was obtained from quantitative phase images by multiplying the phase measurement by wavelength and dividing by 2π. Optical path length is then integrated over an area of interest to obtain Optical Volume. Under the assumption of homogenous refractive index, this metric is proportional to the mass of the cell and can be scaled directly to calculate the dry mass of the region. Dry mass is a common measure used to quantify the content of microscopic objects through phase microscopy, but crucially OV does not rely on selection of this scaling factor.

## Results

To analyze the dual modal system’s spatial and temporal capabilities, simultaneous QPI and FRET images were captured for each cell in the population (N = 29) over a 300 second acquisition period. After an initial 25 seconds of acquisition, a hypo-osmotic stimulus was applied via a syringe-pump at 6mL/min (Fig 2B). Fig 4 shows paired QPI and FRET images of an A431 cell throughout the image acquisition period. Figs 4A and 4D show corresponding QPI and FRET images for the cell prior to hypo-osmotic stimulus. Note the FRET signal’s presence throughout the acquisition due to sustained levels in intracellular Ca^2+^. Fig 4B depicts the cell following the hypo-osmotic stimulus, wherein the measured optical path length of the innermost region has substantially decreased, while the periphery extends in all directions, consistent with the hypothesized flattened profile. The corresponding FRET acquisition (Fig 4E) reflects a similar shape change and a decreased FRET signal. The final paired QPI and FRET images (Fig 4C and 4F) indicate further cell flattening, with decreased localized phase and FRET signals.

**Fig 4.**
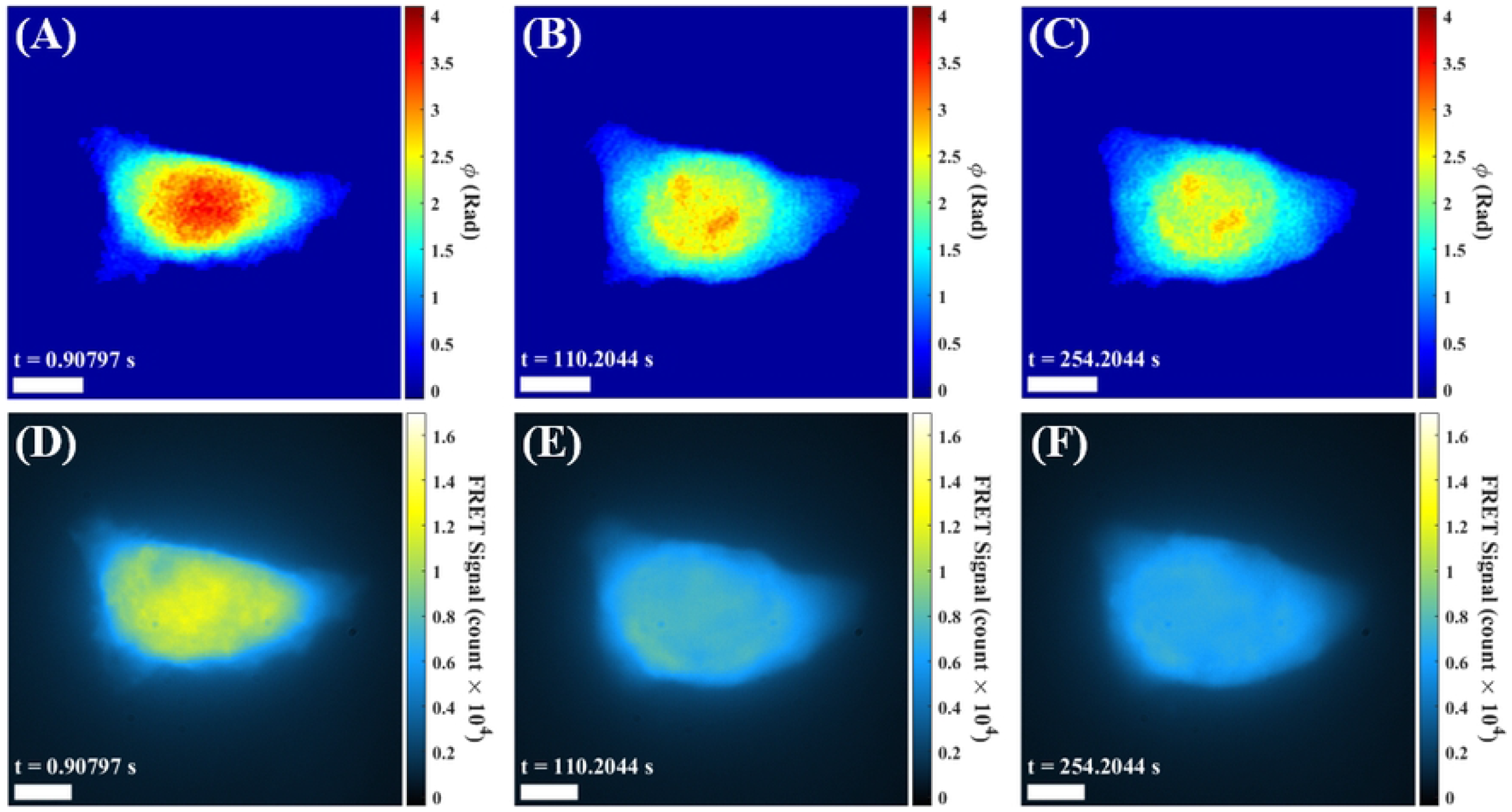
Corresponding QPI and FRET Channel images across the image acquisition period. All scale bars are 10 microns.

To further visualize temporal changes in the QPI signal, image subtraction was used to generate a differential image relative to the initial QPI image frame. Throughout the entire acquisition period, fluid flow was applied to a cell; Fig 5A serves as a control depicting whole cell initial prior to the applied hypo-osmotic stimulus (t = 25 s). The initial response of the cell is captured in Fig 5B, showing a slight overall increase in phase (∼0.3 radians). This increase in phase upon stimulus, could be attributed to mass redistribution stimulated via the influx of Ca^2+^ ions. At approximately 66 seconds (41 seconds after stimulus), Fig 5C depicts the onset of mass redistribution towards the cell periphery. Imaging at longer time scales beyond initial stimulus reveals a damped oscillatory effect between the innermost region and periphery (Figs 5D-F).

**Fig 5.**
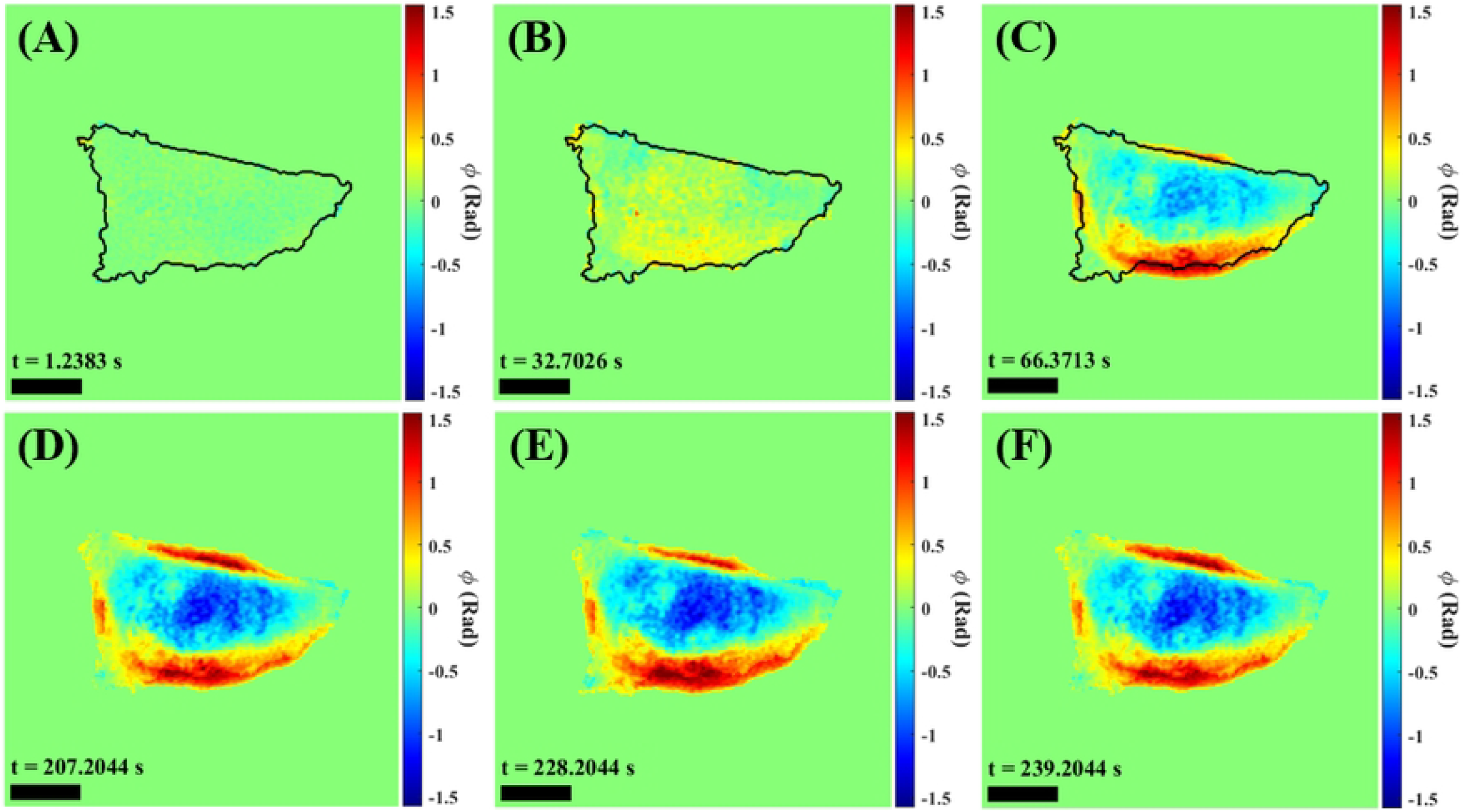
QPI Difference Images, visualizing cellular reorganization and mass transport after hypo-osmotic shock. (A) Cell prior to stimulus. (B) Cell directly after stimulus. (C) Cell approximately 41 seconds after stimulus and beginning to stabilize. (D-F) Cellular mass oscillations can be observed qualitatively at long time scales after stimulus, with mass moving back and forth between the center of the cell and the outer regions. All scale bars are 10 microns.

To further examine regional effects of hypo-osmotic stimulus, automated QPI image segmentation was applied throughout the population. Fig 6A illustrates these phase “layers” applied to a cell of interest. As previously mentioned, phase imaging lacks subcellular structural specificity and automated QPI image segmentation aims to address this drawback. These phase-based layers roughly represent subcellular regions of interest with the inner, middle, and outer layers corresponding to the nucleus, endoplasmic reticulum, and cytosolic periphery, respectively [22]. Moreover, these segmentation layers can be applied to co-registered FRET images to locally analyze the effect of stimuli in a physiologically based manner, thereby overcoming the inability of fluorescent imaging to quantify a relationship between pixel intensity and a corresponding physical region of the cell. Fig 6B shows the registration of a FRET image (red) to QPI image (cyan). As previously mentioned, through this novel dual modality image registration, QPI segmentation layers can be applied to a FRET image for a physiologically relevant regional analysis (Fig 6C). To note, the FRET signal extends beyond the boundary of the QPI image due to the 0.2 radian threshold used in the processing of the phase images.

**Fig 6.**
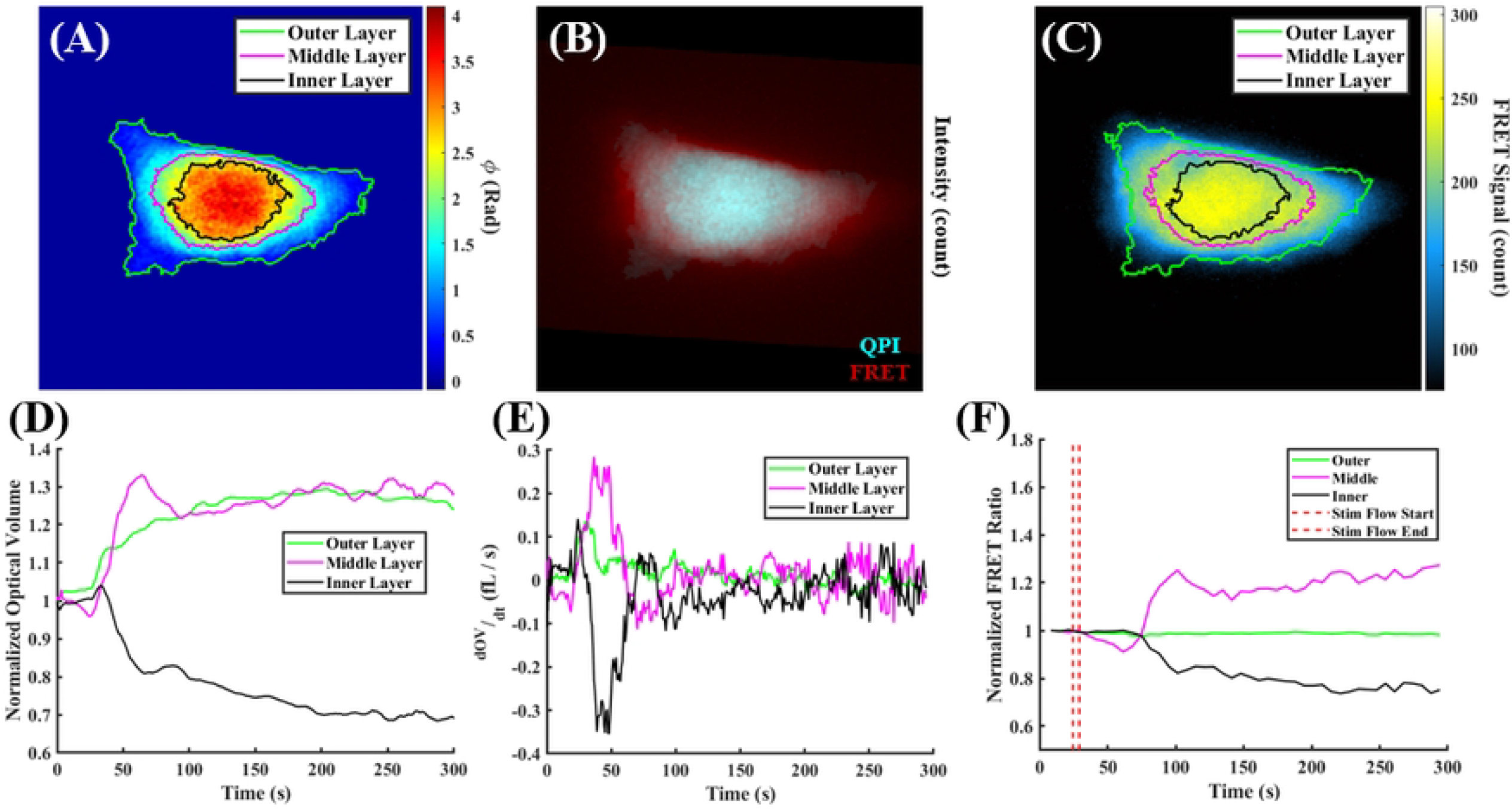
Single cell imaging example of QPI segmentation and QPI-FRET registration to yield regional information (n=1). (A) Visualization of QPI segmentation analysis with inner, middle, and outer layers shown. (B) Registered QPI and FRET image frames of the cell. (C) QPI layers analysis applied to FRET image (reduced FRET count due to image compression). (D) Normalized optical volume for each layer as determined by QPI segmentation. (E) Time derivative of optical volume for each layer. (F) FRET Ratio signal for each registered QPI layer

Using previously described methods, Optical Volume (OV) was calculated from the QPI information and regionally discriminated via the segmentation algorithm. Fig 6D plots normalized Optical Volume of each layer throughout the 300 second acquisition period for a cell of interest. The inner nuclear layer experienced a modest optical volume increase of 4% upon stimulus, followed by a sharp decrease, ultimately falling to 68% of the pre-stimulus measurement. The middle layer, which hypothetically contains the endoplasmic reticulum, increased in OV by 33% following stimulus, wherein the peak OV value of the middle layer succeeds that of the inner layer in the imaging sequence. Subsequently this increase is followed by an approximate 11% decrease and nominal behavior through the remainder of the acquisition. Similarly, the outer layer OV increased upon stimulus (14%) and gradually increased throughout the acquisition, reaching a maximum value of approximately 129% of pre-stimulus optical volume. Interestingly, the mass distribution of the middle layer seems to present oscillatory behavior at long time scales after stimulus was applied. To further analyze the behavior of the segmented layers, the time derivative of the optical volume was calculated and plotted (Fig 6E). The time derivative displays clear local maxima near the application of hypo-osmotic stimulus, allowing for more precise time localization of each layer’s initial response to stimulus. Fig 6E indicates that the inner layer responds first to stimulus, followed by the outer and middle layers. Again, oscillatory behavior is seen in the middle layer. Intriguingly, the middle layer time derivative OV signal appears anti-phase with the inner layer signal. This could potentially suggest oscillatory mass redistributions between the two regions long after stimulus is applied.

Finally, to further investigate overall mass redistribution and analyze whole cell Ca^2+^ ion dynamics after stimulus, A431 cells transfected with Twitch-3 were analyzed using the combined QPI-FRET system (n = 29). Fig 7A depicts averages of the FRET ratio (cyan) and whole cell optical volume (blue) across the population of imaged cells. Additionally, timing of the stimulus is indicated with red dashed lines. Note the FRET signal displays photobleached behavior but appears to temporarily plateau following stimulus.

**Fig 7.**
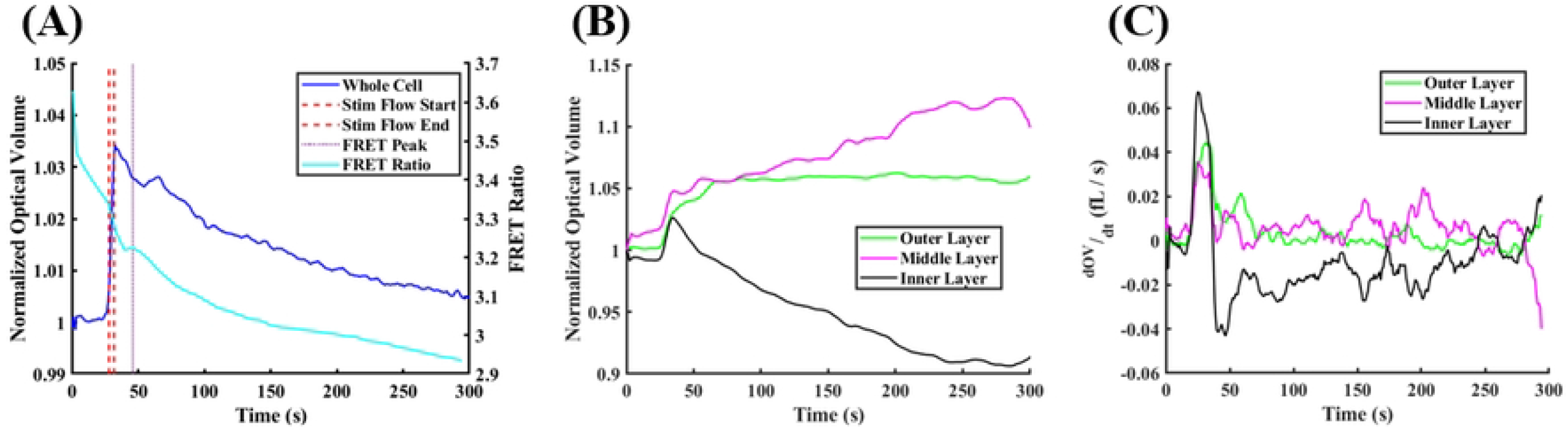
Normalized optical volume, FRET Ratio signal, and Optical Volume time derivative of for cell population (N = 29). (A) Normalized whole cell optical volume and FRET signal over time for the population. (B) Normalized optical volume of cell layers for the cell population over time using our novel QPI segmentation. (C) Time derivative of optical volume of cell layers for the cell population over time using QPI segmentation.

Using optical volume as the preferred QPI metric, population level mass transport dynamics can be summarized. Fig 7A details whole cell optical volume responses across the population, wherein the average whole cell OV increased an average of 3.3% after the application of hypo-osmotic stimulus. This may indicate that the influx of Ca^2+^ ions trigger immediate whole cell reorganization, which can be perceived as a general increase in the phase delay of the cell. Additionally, a local maxima in the Twitch-3 FRET sensor is seen approximately 13 seconds after the spike in whole cell optical volume. Corresponding to an increase in Ca^2+^ ion concentration. Clearly, a distinct time delay occurs between the initial response of cellular reorganization, visualized through optical volume, and the perceived influx of Ca^2+^ ions, visualized through the FRET signal.

Furthermore, populational optical volume information was segmented into a layered analysis. Significantly, different trends are observed for each of the layers (Fig 7B). The inner layer exhibits a sharp increase of approximately 3% in mean optical volume before decaying sharply, eventually reaching a minimum of approximately 91% of the pre-stimulus value. In contrast, the middle layer displays a similarly sharp increase directly following stimulus but rises to 112% of the pre-stimulus value over a significantly longer duration. The outer layer OV moderately rises before stabilizing between 105% and 107% of pre-stimulus optical volume. Populationally, each layer’s behavior remains consistent with the hypothesized whole cell response to stimulus.

As discussed above, an oscillatory effect was observed for single cell data at long-time scales after stimulus (Figs 5D-F and 6D-E). The population OV seems to exhibit a similar trend. Again, the OV time derivative was calculated as shown in Fig 7C. Across the population, the inner and middle layers peak in the time derivative at approximately 24.9 seconds, followed by the outer layer at 33.5 seconds. Comparatively, the inner layer spike is the largest of the three layers, nearly doubling the magnitude of the spike for the middle and outer layers. Furthermore, stimulus response behavior differs across the three layers. Following the spike in OV time derivative, the inner layer experiences a substantial decrease, reaching a negative rate of change nearly equivalent to its initial positive increase. Starkly, the inner layer derivative remains primarily negative while the middle and outer layer derivatives are predominantly positive. Surprisingly, the populational rates of optical volume change for the inner and middle layers appear to oscillate in a mirrored fashion. This trend was observed within isolated cells of interest (Fig 5E), but the trend appears to survive at the population level as well.

## Discussion

During experiments, the combined QPI-FRET optical system facilitated the analysis of cellular dynamics following hypo-osmotic shock in A431 epidermoid carcinoma cells. Moreover, the automated QPI segmentation algorithm introduced here elucidates differential responses to stimulus in the segmented layers. Given these localized responses we hypothesize the height of distinct subcellular regions within the flow channel strongly influence layer-specific responses. The innermost layer, which corresponds to the thicker nuclear region, extends further into the channel compared to the layers. This extension would mean the inner layer experiences the highest magnitude flow and undergoes the largest shear stress (Fig 2B).

The gradational force due to variation in flow rate across the channel produces an influx of Ca^2+^ ions initially localized to the cell center, influencing mass redistribution as seen by an increase of inner layer OV, followed by translation towards the peripheral layers Fig 6F could imply a diffusion of Ca^2+^ ions between the segmented inner and middle cell layers as the increase in one is offset by decrease in the latter. Additionally, given the qualitative change in cell shape as well as the quantitative increase in whole cell optical volume, we can infer that cellular reorganization occurs. The segmentation analysis shows that stimulus results in a flattening of the cell (Fig 2B), and the effect is seen quantitatively across the cell population (N = 29) as observed in Fig 7B. Upon stimulus each layer experiences an increase in optical volume, followed by a distinct layer-specific response that is consistent across the population. Shaked et al. (2010) used QPI to observe a similar increase in whole cell dry mass of chondrocytes experiencing hypo-osmotic shock, reinforcing these findings, as optical volume is directly related to dry mass [23]. Though crucially, the experiments detailed here were conducted over longer time sequences, therefore capturing longer time-scale cell dynamics in addition to the initial stimuli response.

After stimulus cessation, the OV of the inner layer decreased sharply until stabilizing, while the peripheral layers’ OV increased. The innermost region was selected to approximate a nuclear region, and the observed behavior suggests that it behaves distinctly from the rest of the cell body in a hypo-osmotic environment. Previous results by Finan et al. (2009) used light scattering techniques to record different volumetric responses to hypo-osmotic stress between chondrocyte nuclei and the peripheral cell body [24]. While our combined FRET-QPI employs distinctly different contrast mechanisms, these results validate the unique osmotic response properties of the nucleus.

The time derivative of OV for each layer proved an effective tool in highlighting differences among the layers’ distinct behavior at various time scales relative to stimulus. The derivative of the inner layer OV indicates a well-defined increase and decrease after stimulus; however, such a response is not seen in the middle and outer layers. The OV time derivatives across these layers provide a quantitative perspective on cellular morphological response, likely indicating intralayer mass transport in response to stimulus. Specifically, the observed mirrored oscillations in the OV time derivative of the inner and middle layers illustrate this proposed observation of mass transport. The presence of intralayer transport implies signaling that remains activated above basal levels long after applied stimulus. Interestingly, research in chondrocytes has found strong secondary Ca^2+^ peaks that were termed oscillations and occurred approximately two minutes after the initial peak [25]. Oscillatory behavior in the time derivative plots occurs at a similar time scale of about two minutes after stimulus. While it must be noted that cells used here are different than the chondrocytes used in the mentioned study, this may suggest a general trend in cell response, and understanding these oscillations is a topic of future investigation.

Individually, FRET and QPI contribute unique imaging capabilities that advance understanding in cell biology studies. Namely, QPI visualizes cells in a label-free manner and provides morphological information, while FRET sensor engineering provides nuanced reporting of intracellular processes. Here we have presented a novel combination of the two methods that utilize each of their strengths to obtain registered information on mechanical and electrochemical cellular dynamics. The automated segmentation algorithm shows an effective means to address a weakness of QPI, in that individual cellular structures cannot be identified visually. With the segmentation approach, properties of various cellular compartments can be analyzed. Alternatively, recent work in the field has also accomplished this through a machine learning technique [26]. While this approach enables correspondence with fluorescent labels to be established, QPI does not currently provide the same functional information as the Twitch-3 FRET sensor. QPI segmentation analysis, registered to simultaneously captured FRET data, allows for a cell compartment specific measurement of morphological structure and molecular function. Future work will involve further investigation into nuanced cellular processes, with continual advancements in FRET sensor engineering resulting in novel dual modality experimentation. Ultimately, this study demonstrates the capability of a novel dual modality system to dynamically examine live cells tracking their morphological and molecular changes.

## Acknowledgements

This material is based upon work supported by, or in part by, the U. S. Army Research Laboratory and the U. S. Army Research Office under contract/grant number W911NF1910306.

